# Within-host adaptation alters priority effects within the phyllosphere microbiome

**DOI:** 10.1101/2022.08.19.504580

**Authors:** Reena Debray, Asa Conover, Xuening Zhang, Emily A. Dewald-Wang, Britt Koskella

## Abstract

To predict microbiome composition and function over time, it is essential to understand how evolution alters priority effects between resident and invading species. In an experimental evolution study on tomato plants, an early-colonizing bacterial species rapidly evolved to invade a new niche, altering its ecological interactions with other members of the plant microbiome, as well as its effect on the host. Prevailing models have assumed that adaptation primarily improves the efficiency of resident species within their existing niches, yet we conclude that host habitats can offer alternative evolutionary opportunities, complicating the application of existing theory to the community ecology of microbiomes.

## Main

Recent research has revealed a critical role for priority effects, an ecological process whereby community assembly trajectories depend on the order of species arrival, in the assembly, stability, and function of host-associated microbiomes^1,2^. Understanding how resident microbial communities resist and/or facilitate invasions by arriving species is useful for many avenues that include guiding the assembly of healthy microbiomes, designing probiotics that successfully establish within existing communities, and promoting microbiome-mediated resistance to pathogens. Yet we currently have little understanding of how microbiomes evolve within hosts, or how within-host selection alters interactions between resident species and invading species. This is in part because the ecological theories used to understand microbial communities are frequently adapted from other study organisms with slower evolutionary rates.

Prevailing models assume that microorganisms which colonize hosts early in succession evolve to occupy their niches more efficiently, providing an additional competitive advantage against invaders^3^. Early-arriving species would thus resist invasion through both ecological and evolutionary processes, implying that coexistence of competing species should be low, and that microbiomes should be highly resistant to change after initial establishment. Known as the community monopolization hypothesis, this model has received support from several studies of microbial populations evolving in lab culture^4–6^. Yet observations of microbiomes in nature often reveal continuous replacement among strains and species^7,8^, questioning the extent to which adaptation can increase colonization resistance within hosts.

The phyllosphere, or above-ground plant tissues, provides many advantages for studying microbiome assembly. This plant compartment is clearly delineated from the surrounding environment, supports a diversity of microbial species (the majority of which are culturable in the lab), and plays an important role both in individual plant fitness and global nutrient cycling^9^. To interrogate priority effects in this system, we examined the adaptive potential of the early-colonizing bacterium *Pantoea dispersa*. This species comprises a large proportion of the tomato seed microbiome^10^, suggesting that it experiences ample opportunity in nature to colonize new seedlings, and it is one of the most abundant organisms in the phyllosphere of adult tomato plants. However, the relationship between *Pa. dispersa* and its host organisms may be context- or strain-dependent. It can promote plant growth and protect against pathogens^10,11^, yet has also been reported as an opportunistic pathogen of both plants and animals^12^.

We first asked whether priority effects might play an important role in phyllosphere microbiome assembly by introducing *Pa. dispersa* to tomato seedlings either before, simultaneously with, or after a competitor. To capture a range of expected overlap in niches and potential outcomes on host plant phenotype, we selected the following competitors: i) The same species *Pa. dispersa* distinguished by a different selective marker, ii) The plant-protective bacterium *Pseudomonas protegens*, iii) The endophytic plant pathogen *Pseudomonas syringae*. In the first two cases, the eventual community composition strongly depended on arrival order, indicative of priority effects. In contrast, *Pa. dispersa* and *Ps. syringae* reached similar proportions regardless of their arrival order, possibly reflecting differences in their life cycles and spatial localization in plant tissue (Figure **S1**). As arrival order is inevitably correlated with the amount of time between inoculation and harvest in such experiments, it is possible that early arrival appears advantageous only because of population growth, independent of any interactions with other strains. We excluded this possibility by measuring the growth of each strain on plants in the absence of competition, which indicated that the duration of the experiment was sufficient for all strains to reach carrying capacity (Figure **S2**).

To select for increased host association, we repeatedly inoculated tomato seedlings grown in sterile microcosms with replicate populations of *Pa. dispersa* and allowed the bacteria to associate with the plant for one week. Bacteria that successfully colonized the plant were harvested and used to inoculate the next generation of seedlings (Figure **1A**). Following the experimental evolution, we asked whether within-host selection had altered the ability of *Pa. dispersa* to resist invasion and/or invade established populations of competitors (Figure **1B**).

**Figure 1.**
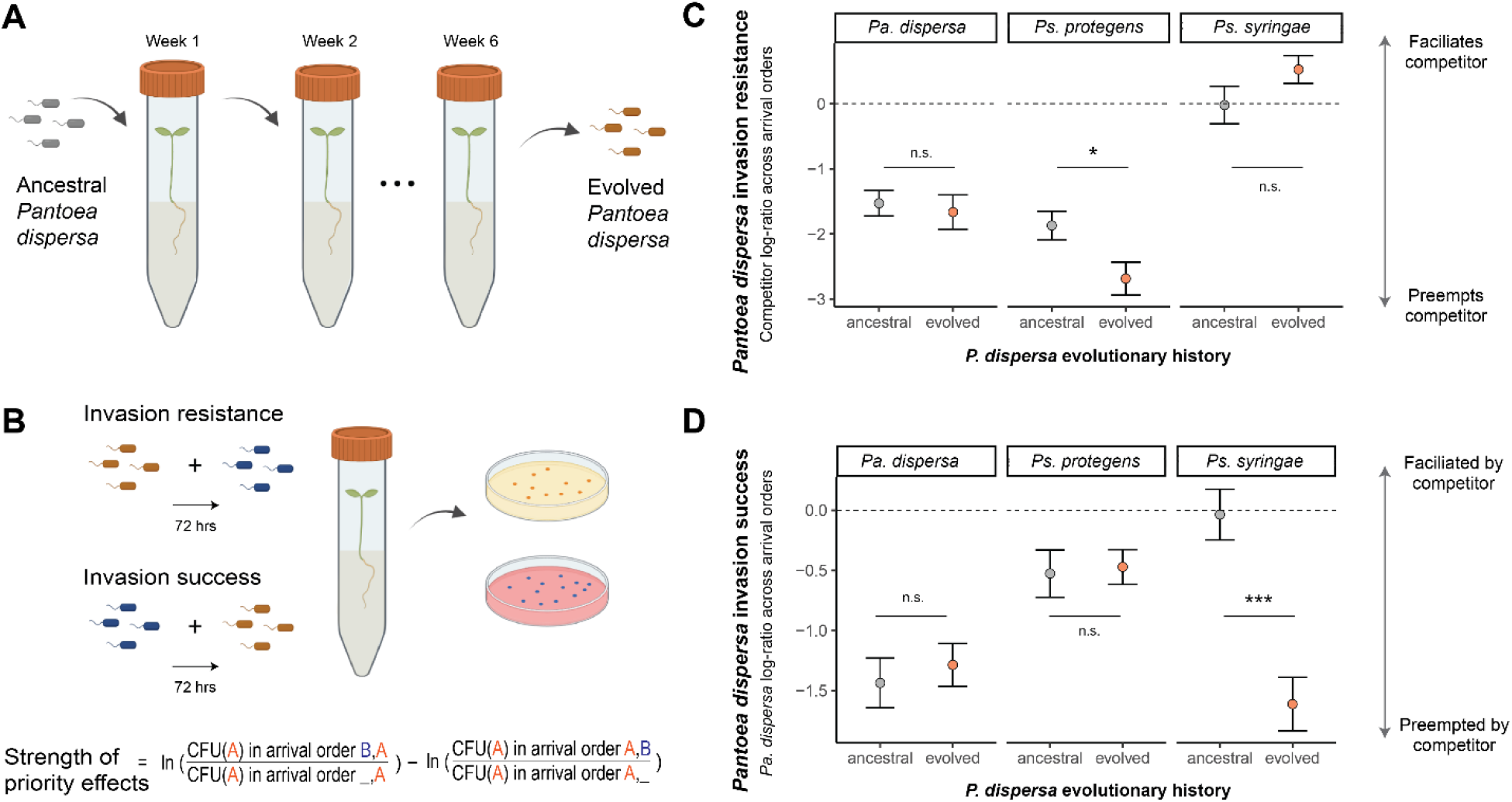
Within-host evolution of the early colonizer *Pantoea dispersa* alters priority effects among bacterial strains. **(a) Schematic of experimental evolution protocol**. Six replicate populations of *Pa. dispersa* were each inoculated onto tomato seedlings via flooding. Seedlings were incubated for 1 week, then harvested to isolate bacteria for the next generation of inoculum. **(b) Schematic of arrival order experiments**. Tomato seedlings were inoculated via flooding with either evolved or ancestral *Pa. dispersa* and a competitor (*Pa. dispersa*, *Pseudomonas protegens*, or *Pseudomonas syringae*), 72 hours apart. Seedlings were harvested another 72 hours after the addition of the second strain and plated on selective media to quantify population sizes. **(c) Resistance to invasion in arrival order experiments**. Evolved populations of *Pa. dispersa* were more resistant to invasion by *Ps. protegens* (t-test, p = 0.018). **(d) Invasion success in arrival order experiments**. Evolved populations of *Pa. dispersa* were less successful at invading *Ps. syringae* (t-test, p < 0.001). Grid panels in (c) and (d) indicate competitor identity and asterisks indicate level of significance: 0.05 < p ≤ 1 (n.s.); 0.01 < p ≤ 0.05 (*); 0.001 < p ≤ 0.01 (**); 0 < p ≤ 0.001 (***). Error bars represent standard error (n=6 replicates per treatment).

After six weeks of experimental evolution within tomato seedlings, populations of *Pa. dispersa* were significantly more resistant to invasion by *Ps. protegens* than their ancestral counterparts, consistent with the hypothesis that within-host adaptation can strengthen priority effects by early colonizers (Figure **1C**). In contrast, evolved populations of *Pa. dispersa* did not improve their ability to invade established populations of competitors (Figure **1D**). Where evolution of invasion success has been reported in the literature, populations were passaged within cultures that had been pre-conditioned by competitors^5^, suggesting that adaptation to the host environment alone may not be sufficient to overcome competition-mediated priority effects. Finally, evolved populations of *Pa. dispersa* were significantly less effective than their progenitors at colonizing plants after the establishment of *Ps. syringae*. This final observation was rather unexpected given that *Ps. syringae*, an endophytic plant pathogen, had a substantially different life cycle than the other strains in the study^13^, and had not previously shown signs of direct competition with *Pa. dispersa*.

To understand this previously unobserved priority effect between *Ps. syringae* and *Pa. dispersa*, we first examined the colonization dynamics of ancestral and evolved *Pa. dispersa* in the absence of competition. The evolved populations appeared to have undergone a life history shift, reaching higher population sizes than their ancestor but only after a period of slow growth (Figure **2A**). Additionally, many plants inoculated with either ancestral or evolved *Pa. dispersa* showed phenotypes reminiscent of disease, such as specks and chlorosis on cotyledons, that were not present in plants inoculated only with sterile buffer (Figure **2B**). To further examine this phenomenon, we adopted a quantitative assay of leaf symptoms previously used to study plant pathogens^10^. We inoculated plants with either ancestral or evolved populations of *Pa. dispersa* and recorded symptom scores on a daily basis for five days, blindly with respect to treatment. This experiment confirmed that evolved *Pa. dispersa* was associated with more pronounced symptom development on leaves (Figure **2C**). We next asked whether it would be possible to leverage priority effects to mitigate the plant phenotypes associated with colonization of the evolved *Pa. dispersa*. When a competing strain of *Pa. dispersa* was introduced three days before the evolved strain, symptom progression was reduced, indicating that priority effects can impact not only community composition but also functional properties (Figure **S3**).

**Figure 2.**
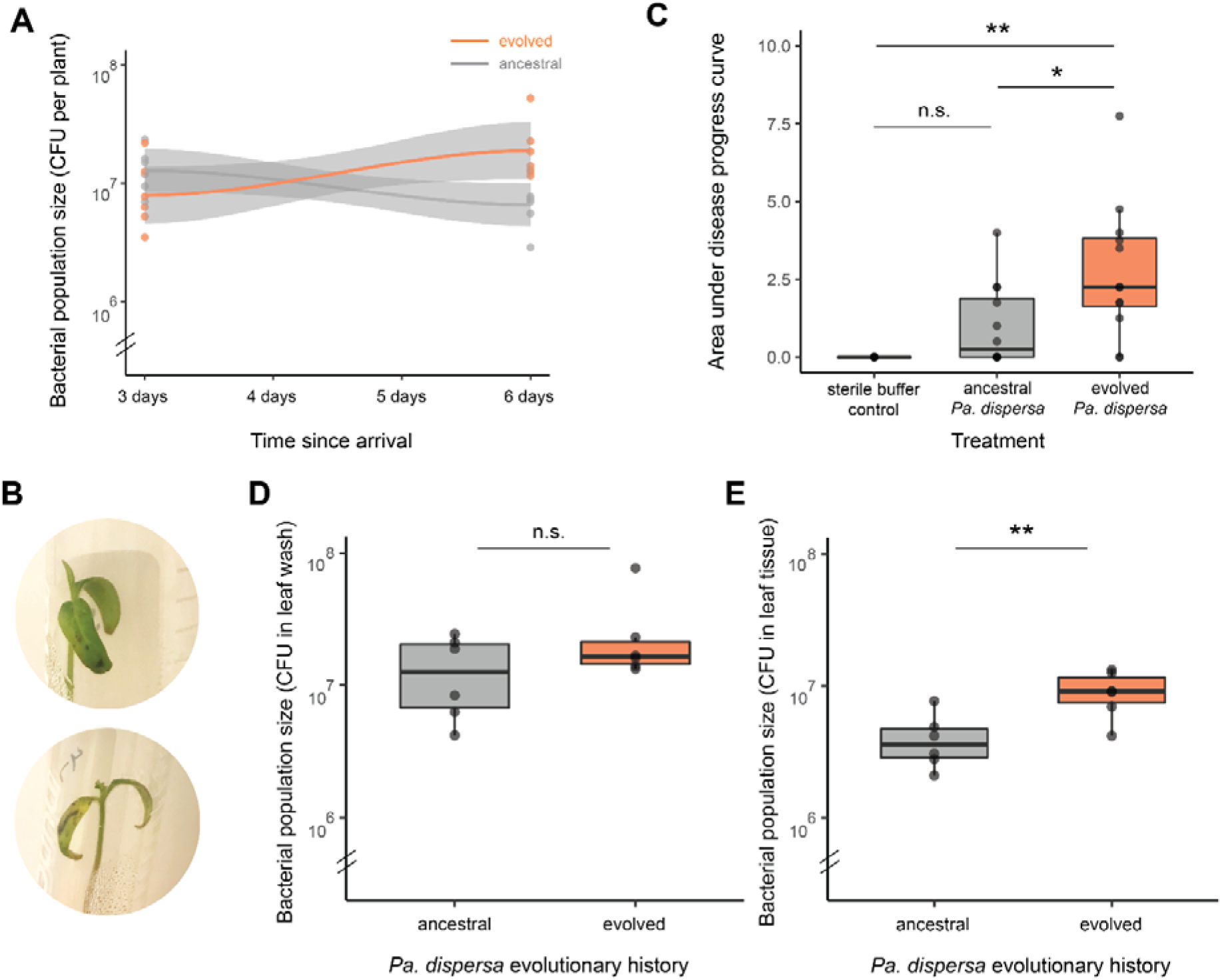
Changes in *Pantoea dispersa* colonization dynamics over experimental evolution. **(a) Life history in the absence of competition**. Population sizes of *Pa. dispersa* after inoculation via flooding. Interaction term between time since arrival and evolutionary history: ANOVA, p = 0.0024. **(b) Images of seedlings inoculated with evolved *Pa. dispersa* populations. (c) Quantitative assay of symptom severity**. A cumulative measure of symptom severity was calculated as the area under the curve of daily symptom scores taken blindly with respect to treatment for 5 days after inoculation. **(d) Bacterial population sizes on leaf surface**. Seedlings were submerged in sterile buffer, sonicated, and vortexed thoroughly. Population sizes in leaf wash were determined via dilution plating. **(e) Bacterial population sizes in leaf tissue**. Following the sonication described in (d), the remaining bacteria associated with seedlings were isolated by homogenizing the seedling in sterile buffer. Asterisks indicate level of significance: 0.05 < p ≤ 1 (n.s.); 0.01 < p ≤ 0.05 (*); 0.001 < p ≤ 0.01 (**); 0 < p ≤ 0.001 (***). Note truncated y-axes and logarithmic scale in panels (a), (d), and (e). Each experiment included one or more seedlings inoculated with sterile buffer and harvested alongside bacteria-treated plants. No bacterial colonies were recovered from these controls.

On the basis of the above observations, we hypothesized that *Pa. dispersa* had evolved to exploit a new niche on the plant. As the colonization success of evolved *Pa. dispersa* depended on whether *Ps. syringae* had previously established, it seemed plausible that *Pa. dispersa* had shifted towards a more endophytic lifestyle. We inoculated plants with either ancestral or evolved *Pa. dispersa* as before, then sampled populations in a spatially resolved manner as follows: First, plants were submerged in sterile buffer and placed in an ultrasonic bath to dislodge bacterial cells from the leaf surface. Then the sonication buffer was removed and plants were homogenized to isolate the remaining bacterial cells associated with the plant. Ancestral and evolved populations of *Pa. dispersa* reached similar population sizes in the leaf wash, but the evolved populations were more abundant within the leaf homogenate (Figure **2D**).

Six weeks of experimental evolution within tomato seedlings considerably altered the temporal and spatial colonization dynamics of *Pa. dispersa* populations. Though these changes were small in magnitude, they were sufficient to affect priority effects among bacterial strains as well as the effect of *Pa. dispersa* on its host. Our findings suggest the evolution of a more intimate association with the plant tissue, possibly mediated by exploitation of plant stomata or improved persistence within other protected sites on the leaf. As in any experimental evolution study, it is important to consider why, if the observed traits are beneficial and evolve readily, the organism has not already acquired them in nature. *Pa. dispersa* appears to be a widely distributed species that has been isolated from plants, animals, and non-host habitats^12,14,15^. It is plausible that improved colonization of tomato seedlings comes with a trade-off at other life stages, on other host species, or in other environments. Another consideration is that the microcosms in this study represent certain conditions (e.g. nutrient limitation, high humidity) that may promote stomatal opening or otherwise make plants particularly vulnerable to exploitation. Indeed, high humidity has previously been shown to promote overgrowth and disease symptoms by endogenous (non-pathogenic) microbiota^16^.

Eco-evolutionary models of community assembly predict that early colonizers should adapt to their hosts and become highly resistant to invasion, yet these predictions are difficult to reconcile with observed fluctuations and replacements in the composition of natural microbiomes. Here, we demonstrate that an early colonizer evolved to exploit a new niche within a short time, changing both the nature and intensity of ecological interactions with other members of the plant microbiome. Our findings suggest that host environments are sufficiently complex and heterogeneous in chemical and physical structure to present possibilities for microbial populations to adapt beyond simply improving their efficiency within their existing niche space. To understand microbiome evolution over host lifespans, it will be critical to further describe the niches available for microorganisms to colonize within hosts, characterize the population sizes and community diversity they can support, and model the expected eco-evolutionary outcomes.

## Methods

### Bacterial strains and selective markers

This study included the following bacterial strains: *Pantoea dispersa* strain ZM1 (originally isolated from tomato plants as reported previously^10^), *Pseudomonas protegens* strain ZDW1 (isolated in this study), and *Pseudomonas syringae* pv. tomato DC3000 (provided by Gail Preston, University of Oxford). The 16S rRNA sequences of all strains are available on the NCBI Sequence Read Archive (BioSample accessions SAMN30116955, SAMN30116956, SAMN30116957). To distinguish competing populations of *Pantoea dispersa* strain ZM1, antibiotic-resistant strains were selected as follows. 500 μL of overnight culture was inoculated into sterile King’s B Medium with either 4 μg/mL rifampicin or 2 μg/mL chloramphenicol. The inoculated culture was incubated overnight at 28°C, and the process was repeated with an increasing antibiotic concentration until resistance was achieved at the final concentration of 20 μg/mL rifampicin or 10 μg/mL chloramphenicol. Strains were then selected for equal competitive ability as their antibiotic-sensitive counterparts *in vitro* by repeatedly co-culturing resistant and sensitive bacteria and measuring the ratios they reached in 24 hours until there were no significant costs of the resistance marker. A single colony of each strain was selected to grow the ancestral stock, which was subsequently used to initiate six experimental evolution populations.

### Arrival order experiments

Money Maker tomato seeds were surface-sterilized in 2.7% bleach (sodium hypochlorite) solution for 20 minutes, then washed three times with 10 mM MgCl_2_ to remove residual bleach. Each seed was placed in a loosely capped, sterile 15 mL tube with 7 mL of 1% water agar. Tubes were covered in foil and maintained in a 21°C chamber until shoot emergence, then moved to a 28°C growth chamber with a 15h day–9h night cycle.

Seedlings were flooded 9-12 days after planting, depending on the experiment. To prepare inocula, overnight cultures were pelleted at 3500 x *g* and washed with 10 mM MgCl_2_ to remove residual media. Each culture was diluted to an optical density (OD_600_) of 0.0015, approximately 10^7^ CFU/mL. The surfactant Silwet L-77 was added at a concentration of 0.015% to facilitate leaf colonization. Tubes were immediately placed on an orbital shaker for 4 minutes, then the inocula were poured off and the seedlings were allowed to dry in a biosafety cabinet. Each flooding trial included one or more seedlings treated with only 10 mM MgCl_2_ and Silwet L-77 to ensure that reagents were sterile.

Three days after the addition of the second species, seedlings were individually weighed and then harvested. Two sterile ceramic beads and 7 mL of 10 mM MgCl_2_ were added to each tube, and tubes were agitated in a FastPrep-24 5G system (MP Biomedicals, Burlingame, CA, USA) at 4 m/s for 60 seconds to homogenize plant tissue. Leaf homogenate was independently diluted twice per sample, and each replicate dilution series was plated on both rifampicin- and chloramphenicol-supplemented King’s B agar plates to distinguish strains in mixed inoculations. Bacterial population sizes were quantified by counting colony-forming units after 2 days of incubation at 28°C.

### Experimental evolution in planta

An overnight culture of *Pantoea dispersa* was pelleted at 3500 x *g* and washed with 10 mM MgCl_2_ to remove residual media. Inocula were prepared by resuspending pellets in 10 mM MgCl_2_ to a bacterial optical density (OD_600_) of 0.0025 and adding 0.015% of the surfactant Silwet L-77 to facilitate leaf colonization. Seedlings were flooded as described above and maintained in the growth chamber for 7 days. At the end of each week, seedlings were collected with sterile forceps into sterile 15-ml Eppendorf tubes and homogenized in sterile 10 mM MgCl_2_ as above. Leaf homogenate was diluted and plated on rifampicin-supplemented King’s B agar plates. For each experimental evolution population, 100 colonies were individually picked from plates, mixed, and grown overnight to generate inocula for the following generation of seedlings.

### Plant symptom quantification

In all cases where symptoms were measured, seedlings were grown in 15 mL tubes and flooded with bacteria at OD_600_ = 0.0015, as described above. Instead of harvesting three days after inoculation to measure bacterial abundances, these seedlings were maintained in the growth chamber to track symptom development on a daily basis. In accordance with previous plant pathology studies, symptoms were scored blindly with respect to treatment using scores that describe different levels of symptom severity^17^. Scores in this study were as follows: no symptoms (level 0), mild speckling (level 1), extensive speckling and/or chlorosis (level 2), and leaf necrosis and/or detachment (level 3). Individual leaflets on the same seedling were scored separately, then combined into an average symptom severity per plant for all analyses and figures to avoid pseudoreplication.

### Statistical analyses

Bacterial population sizes at harvest were determined based on colony counts on selective media. Colony counts of single-species controls were always examined to ensure that no cross-contamination or additional resistance evolution had occurred. We calculated P_ij_, a previously developed metric for quantifying the strength of priority effects^18^, as follows. Here, D(i)_ji_ represents the population size reached by species *i* when it was introduced after species *j*, and D(i)_ij_ represents its population size when *i* was introduced before *j*. D(i)_0i_ and D(i)_i0_ represent the single-species controls of species *i* that were inoculated and sampled at the same time as the corresponding co-colonization treatments.

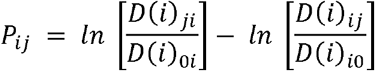

This coefficient takes on negative values if the population size of species *i* is smaller when arriving after *j* than when arriving before; that is, if competition outcomes between *i* and *j* are arrival order-dependent.

We also tried an alternative metric:

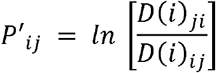

The interpretation of P’_ij_ is the same, but it does not account for the growth of species *i* alone. However, the two metrics were highly correlated and gave qualitatively similar results (Figure **S4**).

Plant symptom progression was analyzed by calculating the area under the disease progression curve (AUDPC), a cumulative measure of symptom severity over time^19^. Differences in cumulative symptom progression over time were assessed using Welch’s t-test that compared the AUDPC across bacterial treatments.

## Acknowledgements

The authors thank Alina Lee and Joy He for assistance with experimental evolution, Callie Chappell for feedback on the manuscript, and members of the Koskella Lab for helpful discussion. RD was supported by the National Science Foundation Graduate Research Fellowship (award number 1650114). The work was supported by the National Science Foundation (award number 1754494).

## Author contributions

RD and BK conceived the study. RD, AC, XZ, and EADW collected the data. RD performed the statistical analyses and drafted the manuscript. All authors contributed to the manuscript revision.

## Competing interests

The authors declare no competing financial interests.

---

**Figure S1.**
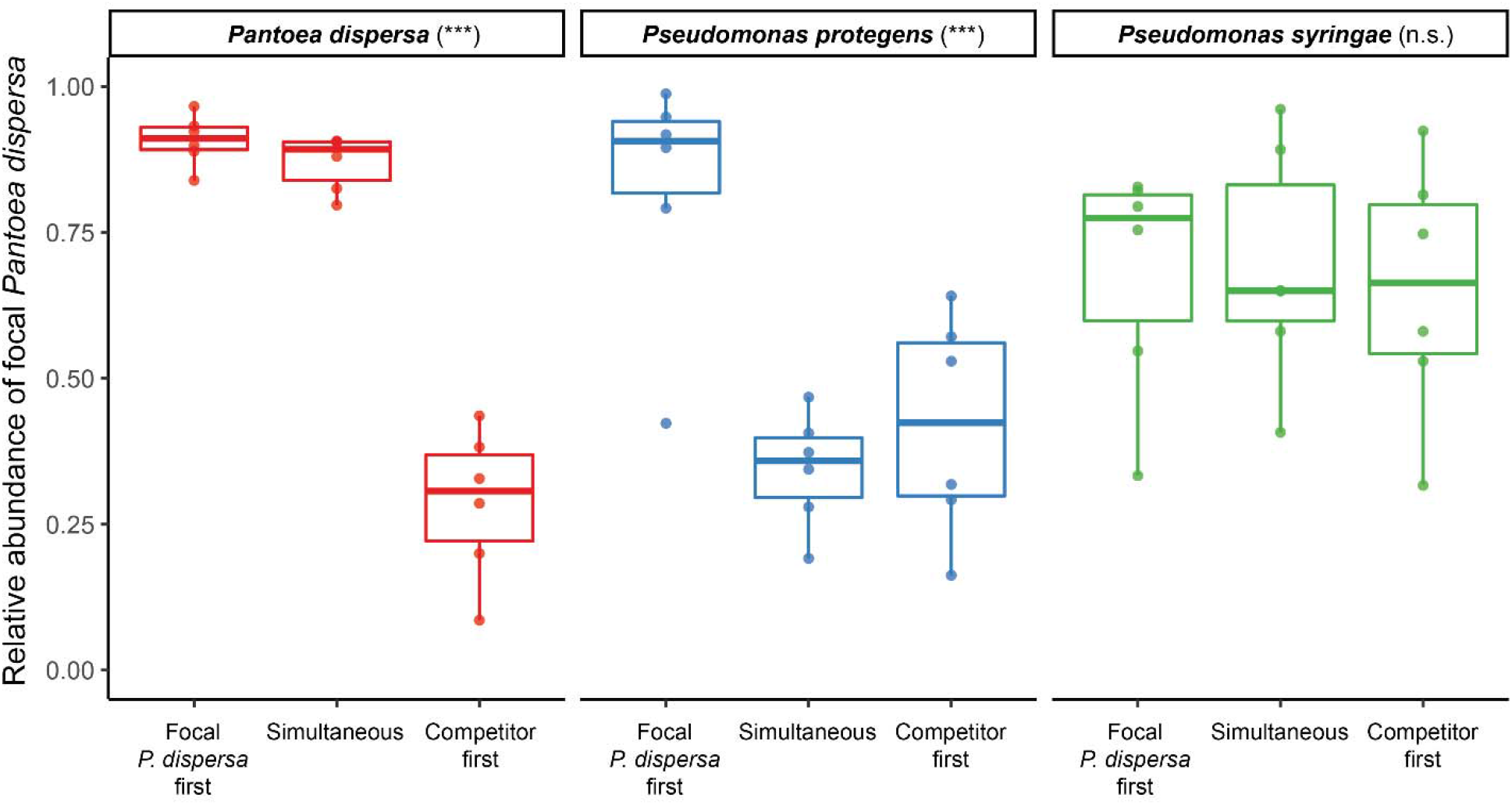
Priority effects among bacteria in the tomato phyllosphere. Relative abundance of *Pantoea dispersa* on tomato seedlings when co-inoculated with either *Pantoea dispersa*, *Pseudomonas protegens*, or *Pseudomonas syringae* across varying arrival orders. Seedlings were flooded either with both strains simultaneously or 72 hours apart. Seedlings were harvested another 72 hours after the addition of the final strain, and homogenized plant tissue was plated on selective media to count colonies. Arrival order significantly impacted strain proportions when both strains were *Pa. dispersa* (ANOVA, F = 107.43, p < 0.001), or when *Pa. dispersa* was co-inoculated with *Ps. protegens* (ANOVA, F = 13.48, p < 0.001), but not when co-inoculated with *Ps. syringae*.

**Figure S2.**
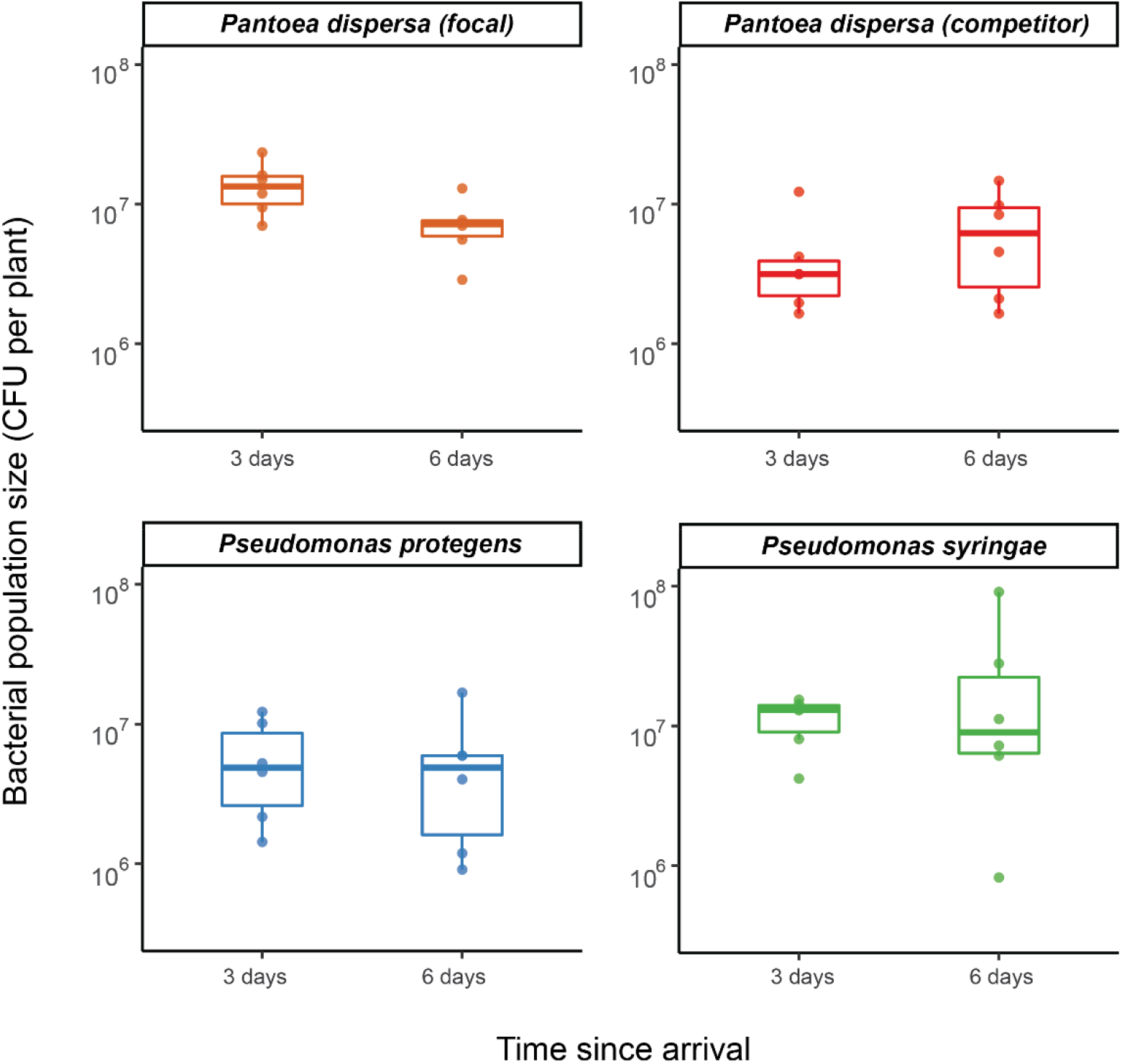
Experimental duration is sufficient to reach carrying capacity on tomato seedlings. Population sizes of *Pantoea dispersa*, *Pseudomonas protegens*, and *Pseudomonas syringae* when inoculated individually onto plants. Seedlings were either harvested directly after 72 hours (as a control for the simultaneous co-inoculation treatments) or flooded with sterile buffer after 72 hours and harvested after 144 hours (as a control for the staggered co-inoculation treatments). At harvest, homogenized plant tissue was plated on selective media to count colonies. None of the strains grew significantly between three and six days after arrival, although the focal strain of *Pa. dispersa* decreased slightly in population size over this time (ANOVA, F = 6.281, *p* = 0.0311).

**Figure S3.**
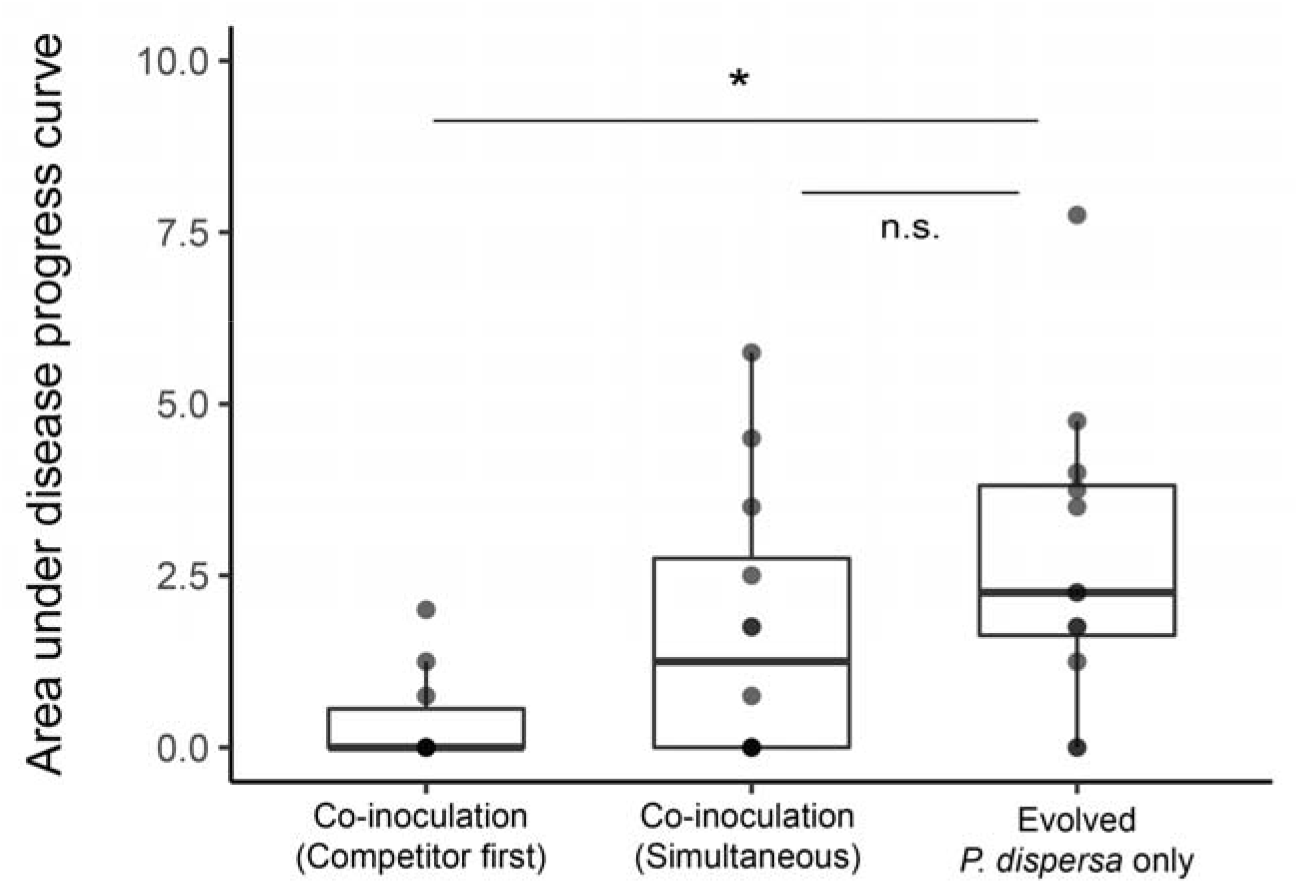
Consequences of priority effects for plant symptom severity. Seedlings were inoculated with evolved populations of *Pantoea dispersa*, which were previously associated with lesions and discoloration of the leaf tissue, and a competing strain of *Pa. dispersa* that does not cause such symptoms. A cumulative measure of symptom severity was calculated as the area under the curve of daily symptom scores taken blindly with respect to treatment for 5 days after inoculation. Each evolved population of *Pa. dispersa* was replicated twice in each treatment category (n = 12 seedlings per treatment), but symptom scores were subsequently averaged for each biological replicate to avoid pseudoreplication (n = 6 per treatment in statistical analysis). Co-inoculation with competing *Pa. dispersa* suppressed symptom development, but only if the competing *Pa. dispersa* arrived first (Tukey’s HSD, p = 0.0242). Asterisks indicate level of significance: 0.05 < p ≤ 1 (n.s.); 0.01 < p ≤ 0.05 (*); 0.001 < p ≤ 0.01 (**); 0 < p ≤ 0.001 (***).

**Figure S4.**
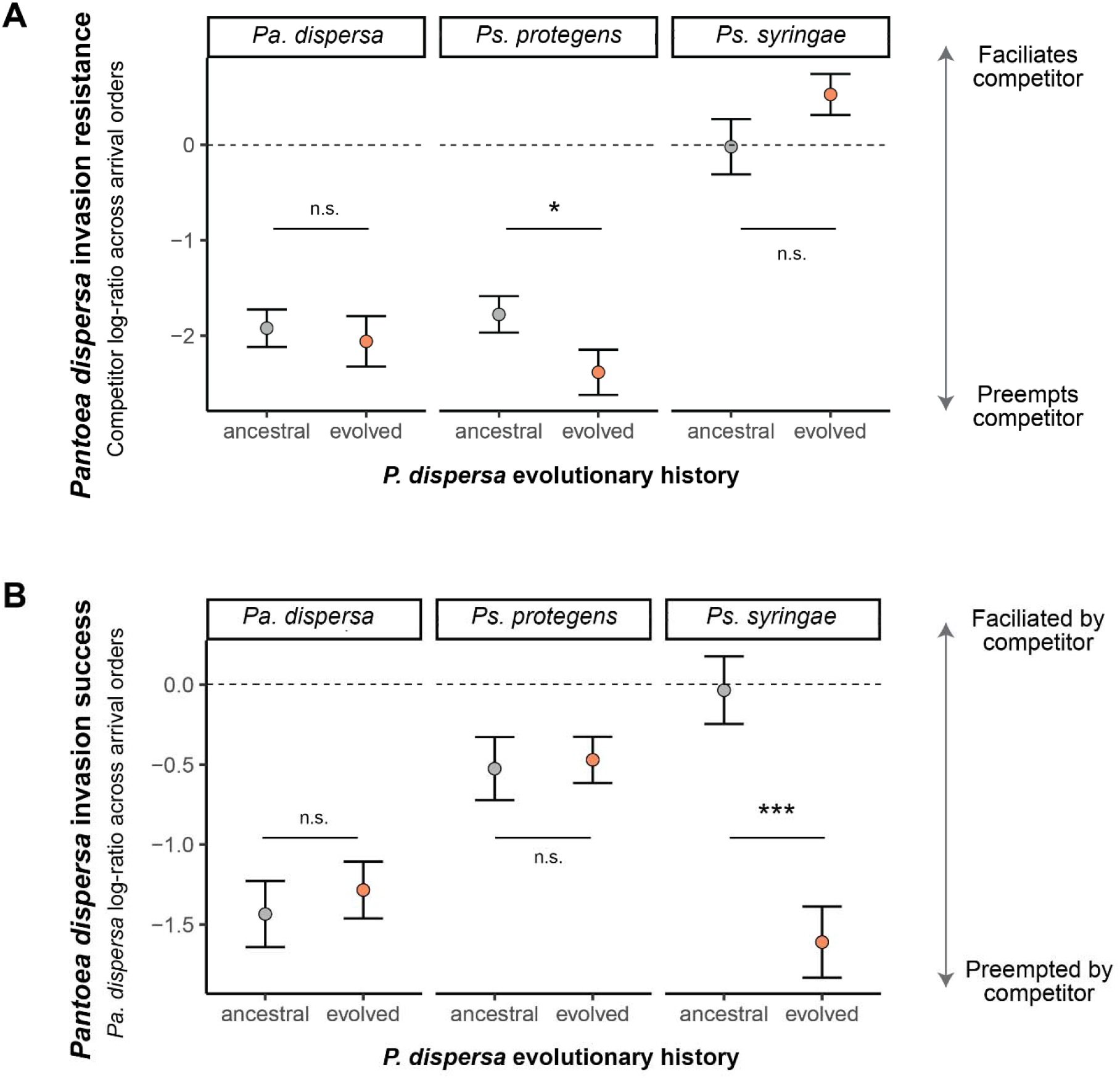
An alternative measure of priority effects recapitulates effects of experimental evolution. **(a) Resistance to invasion in arrival order experiments**. Evolved populations of *Pa. dispersa* were more resistant to invasion by *Ps. protegens* (t-test, p = 0.050). **(d) Invasion success in arrival order experiments**. Evolved populations of *Pa. dispersa* were less successful at invading *Ps. syringae* (t-test, p < 0.001). Grid panels indicate competitor identity and asterisks indicate level of significance: 0.05 < p ≤ 1 (n.s.); 0.01 < p ≤ 0.05 (*); 0.001 < p ≤ 0.01 (**); 0 < p ≤ 0.001 (***). Error bars represent standard error (n=6 replicates per treatment).

## Notes

### Competing Interest Statement

The authors have declared no competing interest.

## References

1. Sprockett, D., Fukami, T. & Relman, D. A. Nat. Rev. Gastroenterol. Hepatol. 15, 197–205 (2018).

2. Debray, R. et al. Nat. Rev. Microbiol. 20, 109–121 (2022).

3. Urban, M. C. & De Meester, L. Proceedings of the Royal Society B: Biological Sciences 276, 4129–4138 (2009).

4. Nadeau, C. P., Farkas, T. E., Makkay, A. M., Thane Papke, R. & Urban, M. C. Proceedings of the Royal Society B: Biological Sciences 288: 20203133 (2021).

5. Chappell, C. R. et al. bioRxiv (2022). doi:10.1101/2022.04.19.487947.

6. Piccardi, P., Alberti, G., Alexander, J. M. & Mitri, S. bioRxiv (2022). doi:10.1101/2022.03.03.482806.

7. Garud, N. R., Good, B. H., Hallatschek, O. & Pollard, K. S. PLoS Biol. 17, e3000102 (2019).

8. Meyer, K. M. et al. ISME J. 16, 1376–1387 (2022).

9. Vorholt, J. A. Nat. Rev. Microbiol. 10, 828–840 (2012).

10. Jiang, L. et al. Sci. Rep. 9, 16354 (2019).

11. Morella, N. M., Zhang, X. & Koskella, B. Phytobiomes Journal 3, 177–190 (2019).

12. Toh, W. K., Loh, P. C. & Wong, H. L. Plant Disease 103, 1764–1764 (2019).

13. Xin, X.-F., Kvitko, B. & He, S. Y. Nat. Rev. Microbiol. 16, 316–328 (2018).

14. Selvakumar, G. et al. World Journal of Microbiology and Biotechnology 24 955–960 (2008).

15. Asai, N. et al. Journal of Medical Case Reports 13, 33 (2019).

16. Xin, X.-F. et al. Nature 539, 524–529 (2016).

17. Rajendran, D. K. et al. Plant Pathol. J. 32, 300–310 (2016).

18. Vannette, R. L. & Fukami, T. Ecol. Lett. 17, 115–124 (2014).

19. Jeger, M. J. & Viljanen-Rollinson, S. L. H. Theoretical and Applied Genetics 102, 32–40 (2001).

